# An EST-SSR based genetic linkage map and identification of QTLs for anthracnose disease resistance in water yam (*Dioscorea alata* L.)

**DOI:** 10.1101/318378

**Authors:** Ranjana Bhattacharjee, Christian O Nwadili, Christopher A Saski, Agre Paterne, Brian E. Scheffler, Joao Augusto, Antonio Lopez-Montes, T. J. Onyeka, P. Lava Kumar, Ranajit Bandyopadhyay

**Affiliations:** International Institute of Tropical Agriculture (IITA), Oyo Road, PMB 5320, Ibadan, Nigeria; Michael Okpara University of Agriculture, Umudike, Abia state, Nigeria; National Root Crops Research Institute, Umudike, Umuahia, Abia State, Nigeria; Institute for Translational Genomics, Genomics and Computational Biology Laboratory, Clemson University, Clemson, SC, USA; Genomics and Bioinformatics Research Unit, USDA-ARS, Stoneville, Mississippi, USA

## Abstract

Water yam (*Dioscorea alata* L.) is one of the most important food yams with wide geographical distribution in tropics. One of the major constraints to water yam production is anthracnose disease caused by a fungus, *Colletotrichum gloesporioides* (Penz.). There are no economically feasible solutions as chemical sprays or cultural practices, such as crop rotation are seldom convenient for smallholder farmers for sustainable control of the disease. Breeding for development of durable genetic resistant varieties is known to offer lasting solution to control endemic disease threats to crop production. However, breeding for resistance to anthracnose has been slow considering the biological constraints related to the heterozygous and vegetative propagation of the crop. The development of saturated linkage maps with high marker density, such as SSRs, followed by identification of QTLs can accelerate the speed and precision of resistance breeding in water yam. A total of 380 EST-SSRs were used to generate a saturated linkage map. About 60.19% of SSRs showed Mendelian segregation pattern, however, it had no effect on the construction of linkage map. All 380 EST-SSRs were mapped into 20 linkage groups covering a total length of 2559.66 cM, which agrees with the diploid nature (2n = 2x = 20) of the parents used in the cross. Majority of the markers were mapped on linkage group 1 comprising of 97 EST-SSRs. This is the first genetic linkage map of water yam constructed using EST-SSRs. QTL localization was based on phenotypic data collected over a 3-year period of inoculating the mapping population with the most virulent strain of *C. gloeosporoides* from West Africa. Using the mean permutation value of LOD scores as threshold value for declaring a putative QTL on all linkage groups, one QTL was consistently observed on linkage group (LG) 14 in all the three years and average score data. This QTL was found at position interval of 71.12 – 84.76 cM explaining 68.94% of the total phenotypic variation in the average score data. The high marker density allowed identification of QTLs and association for anthracnose disease, which could be validated in other mapping populations and used in marker-assisted breeding in *D. alata* improvement programmes.

## Introduction

Yams (*Dioscorea* spp.) are among the primary food crops, and rank second after cassava, in West Africa. More than 54 million tons of yams are produced in sub-Saharan Africa annually on 4.6 m ha. Over 95% of this production comes from the “yam belt” of West Africa that includes Nigeria, Benin, Togo, Ghana, and Côte d’Ivoire [1]. Although yam production in Africa is 40% that of cassava, the value of yam production exceeds all other African staple crops, and is equivalent to the summed value for the top three cereal crops (maize + rice + sorghum) (http://www.iita.org). Yam tubers are the preferred staple food in the region considering its nutritional qualities such as carbohydrates, vitamin C, essential minerals and dietary fiber. The consumer demand for yams in West Africa is very high but its productivity is declining in the region due to several biotic and abiotic constraints.

There are about 600 wild and domesticated *Dioscorea* species. White Guinea yam (*D. rotundata)* is the most important species in the “yam belt” of West and Central Africa. It is indigenous to West Africa, as is the yellow Guinea yam (*D. cayenensis)*. Water yam (*D. alata*) originated in Asia and is the most widely distributed cultivated species in the world [2]. It is superior to most cultivated yam species because of its genetic potential to yield even under low to average soil fertility, ease of propagation through production of bulbils and high multiplication ratio, early vigor for weed suppression, and low post-harvest losses [3]. It is a very important food security crop in many tropical and sub-tropical countries in West Africa, Asia, Latin America and The Pacific, while its production is threatened by the fungal pathogen *Colletotrichum gloeosporoides* (Penz) that causes anthracnose disease [4, 5]. The disease causes mild to acute leaf necrosis and shoot die-back resulting in yield losses of up to 90% during severe conditions [6, 7]. The disease is also considered to be responsible for the disappearance of some of the popular cultivars of *D. alata* in the Caribbean. Recent studies focusing on biodiversity of *C. gloeosporoides* that associates with yam in the French West Indies [8, 9], in Nigeria [10, 4] and West Africa [11] revealed high genetic and pathogenic variance among the isolates. This high genetic and pathogenic variation in the populations of *C. gloeosporiodes* suggests that there is high likelihood for the breakdown of existing resistance and development of new strains of the pathogen [12, 13, 14]. Screening exotic germplasm accessions and selecting for new sources of durable disease resistance is the most reliable approach to long-term management of this disease. However, it is challenging to adequately evaluate yam genotypes for anthracnose disease response under multi-location field conditions because of the high variation among the pathogen isolates and variability in symptoms development across seasons [13, 14]. Furthermore, the standardization of screening protocol for anthracnose disease scoring is underway (Kumar et al., unpublished), although disease screening in controlled environments is favorable for both disease development and plant growth [15, 16]. The genetic improvement of water yam at International Institute of Tropical Agriculture (IITA), Nigeria; Center for Tropical Crops Research Institute (CTCRI), India; Centre de recherche Antilles (INRA), Guadeloupe; and Centre de cooperation international en recherche agronomique pour le développement (CIRAD), Guadeloupe has been ongoing to develop high-yielding anthracnose resistant varieties through classical breeding using phenotypic observations. It is however very difficult to breed for resistant varieties using conventional methods due to several constraints related to the nature of the crop such as long growth cycle (9-11 months), dioecious, poor to erratic flowering, polyloidy, vegetative propagation and heterozygous genetic makeup [17, 18]. The use of molecular techniques, such as the candidate gene approach to dissect the underlying gene (s) for the trait of interest, would accelerate efforts in introgressing disease resistance into preferred genetic background. Some earlier work investigating the genetic inheritance of anthracnose disease in water yam showed that resistance is likely to be dominant and quantitatively inherited [15, 19]. Mignouna et al [20] used AFLP markers to demonstrate that there is a single major dominant locus, designated as *Dcg*-1, controlling resistance in the breeding line, TDa 95/00328 and this resistance is isolate specific. Moreover, it was also suggested that combining both conventional and molecular approaches would be the best approach to develop varieties with a wide range of stable resistant gene (s) to sustain against a broad spectrum of fungal pathogens for yam improvement. It is an already well-established fact that speed, economics, and precision can be improved in breeding cycles by using molecular markers and dense genetic linkage maps. The availability of molecular markers and linkage maps allows identification of locations or markers linked with the trait of interest, thus making it possible to manipulate and understand the inheritance of quantitative traits such as disease resistance. It is also possible to precisely localize quantitative trait loci (QTLs) on the linkage map.

Attempts have been made to construct linkage maps in *Dioscorea* spp. and the first map was developed using AFLP and SSR markers in dioecious diploid wild species *D. tokoro* [21]. Similarly, the AFLP and RAPD-based genetic maps were developed in *D. alata* by Mignouna et al [3] and Petro et al [19]. The SSR markers developed in *D. tokoro* were not suitable for application in cultivated species such as *D. rotundata* and *D. alata* due to their incompatibility [22]. The efforts on development of linkage maps for QTL detection in *D. alata* has been limited due to unavailability of enough molecular markers that are efficient, user-friendly, and co-dominant such as simple sequence repeat (SSRs) markers or single nucleotide polymorphism (SNPs) markers. Subsequently, a total of 44,757 EST-sequences, with an average length of 500 bp, were generated from the cDNA libraries of two resistant (TDa 87-01091 and TDa 95/00328) and one susceptible (TDa 95-310) genotypes of *D. alata* for anthracnose disease [22]. These EST-sequences were further annotated to obtain 1,152 EST-SSRs [23] and a total of 380 polymorphic EST-SSRs were used in the present study to develop the genetic linkage map and identify QTL (s) for anthracnose disease.

## Materials and Methods

### Mapping population

A mapping population of *D. alata* consisting of 94 progenies developed from a cross of two diploid genotypes including TDa 95-310 (susceptible male parent) and TDa 95/00328 (moderately resistant female parent) was used in the study. Both parents are breeding lines developed at IITA and showed differential reactions to several isolates of *Colletotrichum gloeosporioides* [14]. Tubers of 96 yam genotypes, including both parents, were sliced into minisetts of 20-25 g and planted in plastic pots filled with sterilized soil. The experiment was carried out in the greenhouse for three months, an optimal age for vine cutting, and the vines were cut from each genotype to establish a new set of plants of uniform age and growth across all the genotypes before inoculation with the most virulent isolate of *C. gloeosporioides*, Kog01R1. The mini-tubers from each genotype was harvested which served as the planting materials of the following two years for assessment of anthracnose disease for a total of three years.

### Screening of mapping population for anthracnose disease

In the anthracnose screening, a virulent fast-growing salmon (FGG) isolate of *C. gloeosporioides* (Kog01R1), representing the most virulent group in the population of isolates sampled from five different countries in West Africa (Nigeria, Ghana, Ivory Coast, Togo and Benin) [11], was used. All 96 genotypes were planted using a completely randomized design with four replications and the parents were considered as controlled checks in all three years’ evaluations. Whole-plant inoculation was used for disease-assessment and scoring [25, 26, 13]. All the plants were spray-inoculated with spore suspension of Kog01R1 isolate (10^6^ spores mL ^−1^) in the screenhouse using approximately 1 mL of inoculum per plant. The inoculated plants were scored for anthracnose severity from 3^rd^ day onwards until 11^th^ day on a scale of 1-5 using the percentage whole plant-area scoring method [27] with slight modification according to Onyeka et al [13]. Anthracnose score 1 represented high resistance with a leaf area damage of ≤ 2% and score 5 represented high susceptibility with a leaf area damage of ≥ 50%.

### DNA extraction

Genomic DNA from all 94 individuals and two parents was extracted from young leaf tissues collected before inoculation of the plants according to the procedures described by Sharma et al. [28]. DNA was extracted from 1 g of fresh leaves of 3-months-old screenhouse grown plants. DNA quality was assessed by running the samples on 0.8% agarose gel following electrophoresis, and the quantity was estimated on NanoDrop2000 spectrophotometer, using the ratio of absorbance at 260 nm and 280 nm to assess the concentration in ng/μl and purity level of the DNA.

### EST-SSR analysis

The DNA samples collected from the mapping population and the two parents were shipped to GBRU, USDA-ARS, Stoneville for further analysis. A total of 1152 EST-SSRs were generated from >40,000 EST-sequences [23], which were tested for polymorphism on the two parents. A total of 435 SSRs were polymorphic, 329 did not produce any amplified products, and remaining 388 SSRs were monomorphic. Among 435 polymorphic SSRs, 380 showed good polymorphisms and were bi-allelic while 55 showed multi-allelic banding patterns. Only the bi-allelic 380 SSRs were selected for further analysis and their forward and reverse sequences can be obtained from the corresponding author on request.

Primers were obtained from Integrated DNA Technologies (IDT, Coralville, IA) and normalized at 6 nmols in a 96-well plate. For genotyping, a tailing method was used to add a fluorescently labeled tag to the final PCR product as a third primer homologous to the tail. Forward primers were 5′ tailed with the sequence 5′-CAGTTTTCCCAGTCACGAC-3′ and the third tailed primer was Primer 5′-CAGTTTTCCCAGTCACGAC-3′ labeled with 6-carboxy-fluorescein. The reverse primers were tailed at the 5′ end with the sequence 5′-GTTT-3′ to promote non-template adenylation [29]. The 5 μl PCR reaction contained 10 ng genomic DNA, 0.4 μl Titanium Taq DNA Polymerase (Clontech, Mountain View, CA), 0.5 μl Titanium buffer, 0.04 μl dNTP (25mM), 0.8 μl FAM labeled primer (05 pmol/μl), 0.4 pmol of forward primer and 1.2 pmol reverse primer). The reaction was run on a BioRad thermocycler (Hercules, CA) at 95 °C for 1 min, 60 °C for 1 min (two cycles), 95 °C for 30 s, 60 °C for 30 s, 68 °C for 30 s (27 cycles), and a final extension at 68 °C for 4 min. Fluorescently labeled PCR fragments were analyzed on an ABI 3730XL DNA Analyzer with ROX 500 size standard and data-processed using GeneMapper Version 5.0 (all three from Applied Biosystems, Foster City, CA). Each primer pair was tested on both parents and the data compared to determine the number of primer pairs that amplified, the allele number and size, and their polymorphic rate.

### Linkage map construction and QTL mapping

The EST-SSR data was summarized across all individuals following a typical diploid Mendelian inheritance for an F_2_ population, considering the cross-pollinated heterozygous nature of yam and its maintenance over the years through clonal propagation [3]. Missing data was considered, progenies and markers with high missing data were removed from the data. Least significant difference (LSD) and G X E analyses was performed where by data collected from the various years was considered as environmental factor. Linkage map construction was performed using JoinMap and R/qtl [30] for comparison, with Kosambi map function to convert recombination frequencies to genetic distances in centiMorgan (cM) units and to know the correct position of makers for this population. Markers were assigned to linkage groups using a minimum independence LOD score of 3.5 while regression mapping method was used to build the linkage maps. Heatmap which explained correlation between the different markers on the chromosomes was performed using R/qtl package.

The datasets of the 94 progenies (marker data and anthracnose scoring data) combined with the linkage map were used to identify regions associated with anthracnose disease. Haley-Knott regression method [31] and simple interval mapping (SIM) method with maximum step-size of 2 Centimorgan (cM), permutation of 1000 iteration and threshold method [32] at 1% significance were used for QTL detection. The proportion of the phenotypic variance explained by the marker segregation was determined by the R^2^ value. Statistical test of significance (ANOVA) was used to find significant QTLs and to determine the percentage contribution of each QTL to phenotypic variation. All analysis for QTL identification was carried out using R/qtl [33] and the software Breeding View 3.0.9 [34]. Simple Interval Mapping was also performed for QTL analysis to capture the variation of each year and the average score across all the years. Significant QTL were then mapped using Mapchart 2.3 software for mapping QTLs and the flanking region was created.

## Results

### Evaluation for anthracnose resistance

A set of 41 *C. gloeosporiodes* isolates collected from five different countries in West Africa was studied for their differences in causing anthracnose on different *D. alata* varieties using detached leaf technique and whole-plant inoculation [14]. Eleven out of 41 isolates showed high virulence and a PCA graph was used to estimate their virulence based on the distance from the centre (Fig 1). Three isolates such as Kog01R1 (from Nigeria), Gha09A1 (from Ghana) and Tog12A1 (from Togo) showed high virulence and were grouped into highly virulent (Kog01R1) and virulent (Tog12A1 and Gh09A1) categories. These isolates also caused significant necrosis on the yam plants with Kog01R1 showing maximum virulence in both detached leaf and whole-plant inoculation techniques (Table 1, Fig 2). The highly virulent isolate Kog01R1 from Nigeria was selected to evaluate the mapping population consisting of the two parents and their ninety-four progenies. Kog01R1 has the features to exhibit high virulence at all stages of experiments carried out and it also showed the highest rate of infection in all the tested yam varieties, although there was variation in the rate of infection [14]. It is similar in characteristics to the fast-growing gray (FGG) isolates of *C. gloeosporioides*, and Abang et al [24] stated that FGG group is of high virulence as observed based on morphological characterization of isolates.

**Fig. 1.**
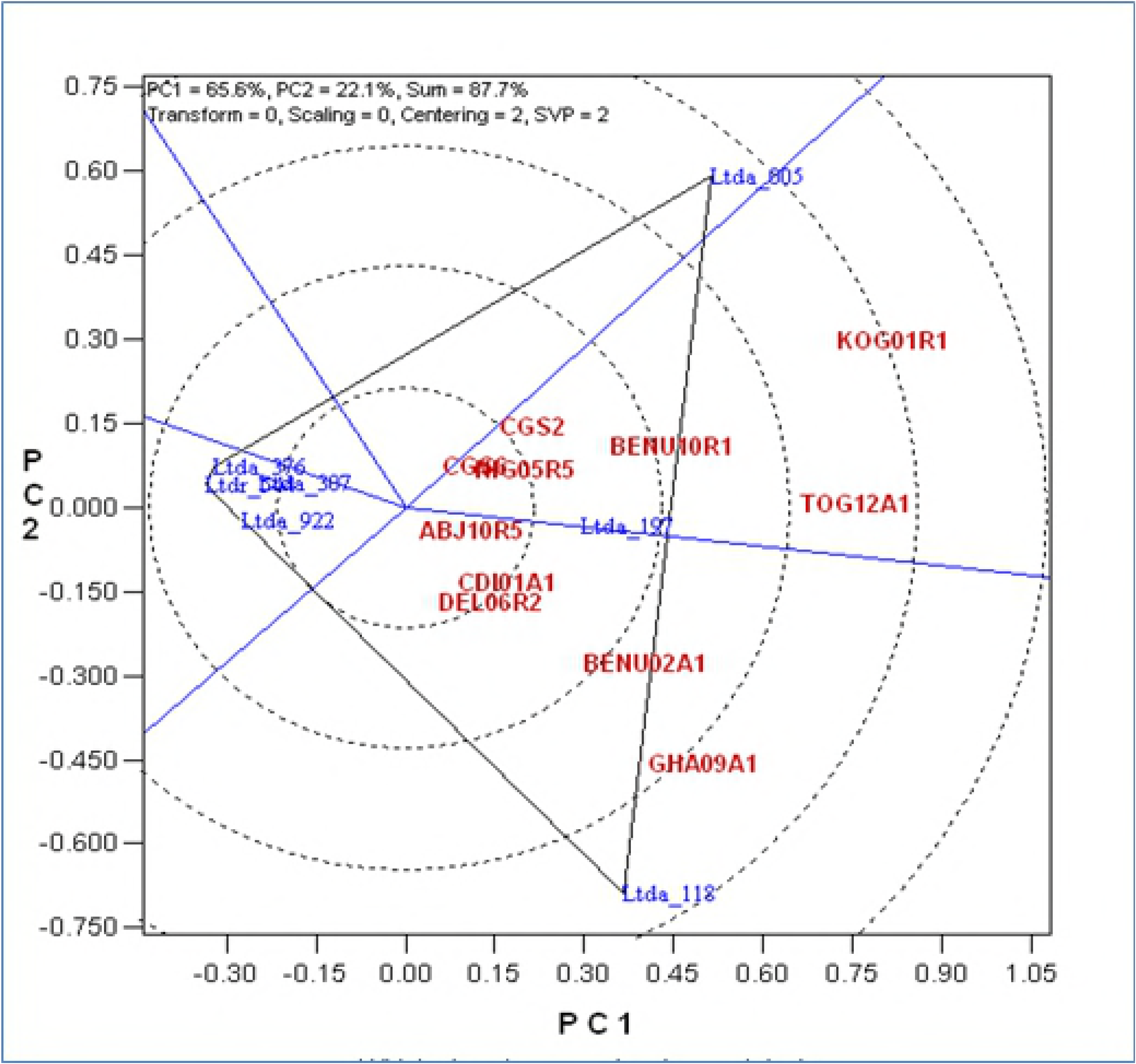
Whole plant assay of eleven out of 41 isolates of *C. gloeosporioides* on different *D. alata* varieties for anthracnose disease. The principle component analysis (PCA) graph indicates the virulence of 11 isolates when tested on different *D. alata* varieties based on whole plant assay.

**Table 1.** Characterization of *C. gloeosporioides* isolates based on anthracnose virulence of the isolates on different yam varieties for detached leaf technique (DLA) and whole plant assay (WPA)

| Isolates | TDa 99/00240 |  | TDa 93–36 |  | TDa 92.2 |  | Tda 95/00005 |  | Virulence |
| --- | --- | --- | --- | --- | --- | --- | --- | --- | --- |
|  | DLA | WPA | DLA | WPA | DLA | WPA | DLA | WPA |  |
| KOG01R1 | 4.7 | 3 | 3.1 | 3.5 | 3.2 | 3 | 2.5 | 3 | Hv |
| GHA09A1 | 4.1 | 3.5 | 3 | 3 | 2.6 | 3 | 2 | 3 | V |
| TOG12A1 | 3.9 | 2 | 2.8 | 3 | 2.4 | 3 | 2 | 2 | V |

DLA: Detached leaf assay; WPA: Whole plant assay; Anthracnose disease scoring on 1-5 scale with 1 = highly resistant; 2 = resistant; 3 = moderately resistant; 4 = susceptible; 5 = highly susceptible

**Fig. 2.**
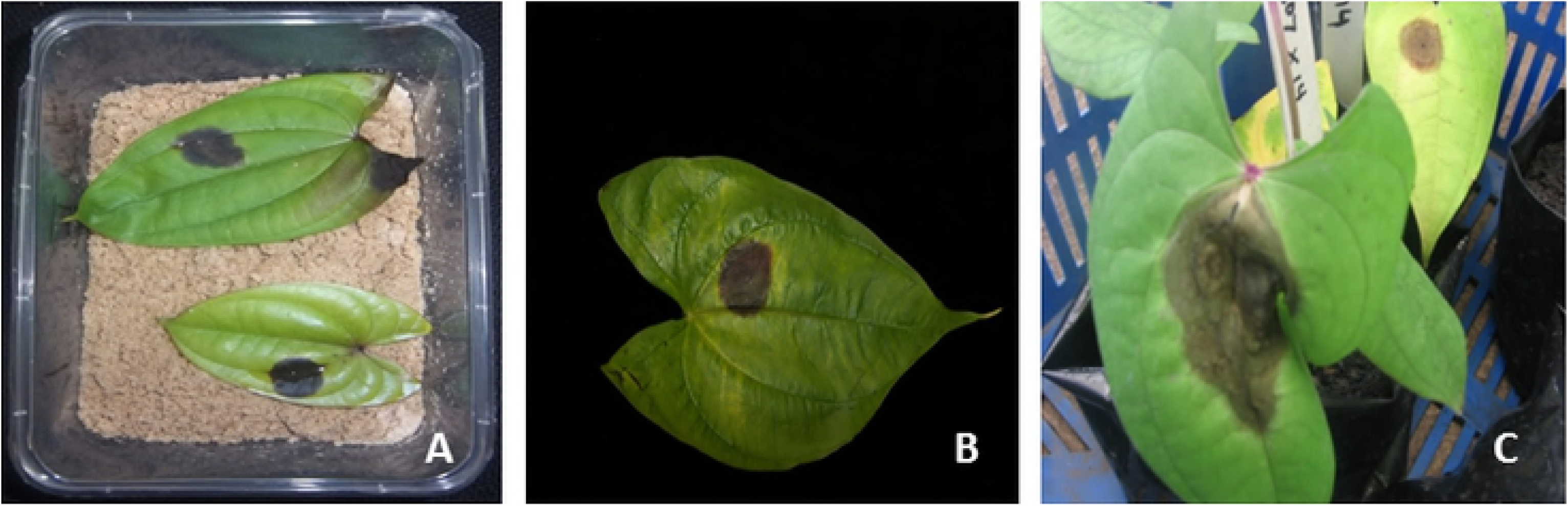
Characterization of *C. gloeosporioides* isolates for virulence using a detached leaf technique and whole plant inoculation. Phenotypic evaluation of genotypes based on detached leaf technique and whole plant inoculation. The disease scoring was based on symptoms and disease progression after 0 DAI, 7 DAI and 15 DAI on a scale of 1-5. DAI: days after inoculation.

The phenotypic evaluation of bi-parental population of 96 genotypes (2 parents and 94 progenies) for anthracnose disease was carried out for three years in greenhouse using whole-plant inoculation technique. The parental clones showed a contrasting level of resistance. TDa 95/00328, moderately resistant female parent, showed a higher resistance to the isolate than the highly susceptible male parent, TDa 95-310. The phenotypic scoring of 94 progenies based on whole plant anthracnose severity resulted in 5.2% of highly resistant, 25.0% resistant, 44.8% moderately resistant, 24.0% susceptible, and 1.0% highly susceptible groups as their mean anthracnose incidence across three years [14] (Fig 3).

**Fig. 3.**
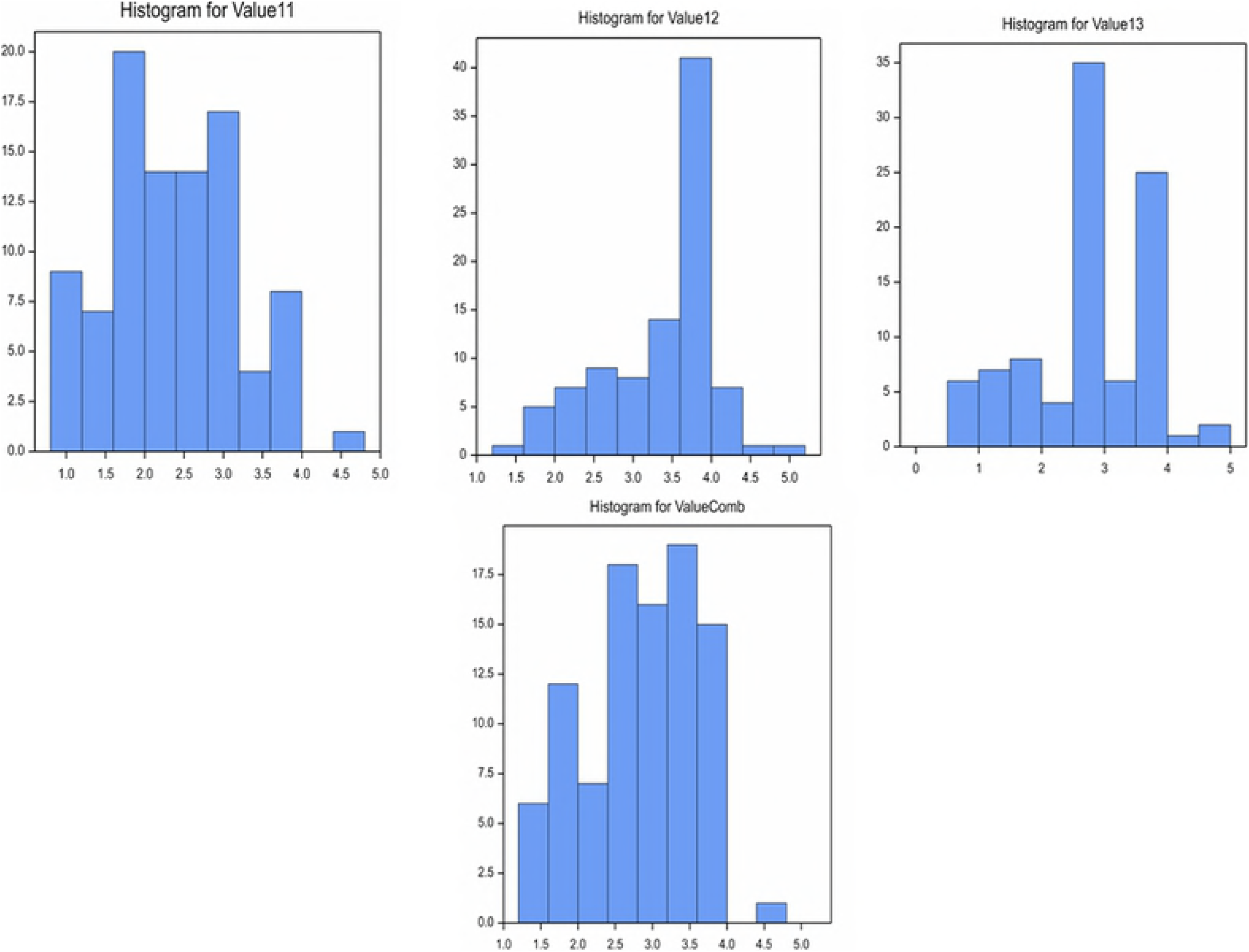
Mean phenotypic distribution of 94 progenies for anthracnose disease for individual years and average across three years. Frequency distribution of 94 progenies based on phenotypic scoring on a scale of 1-5 following whole plant inoculation assay.

### Linkage mapping

The genetic linkage map was constructed using 380 EST-SSR marker data on both parents and 94 segregating progenies (Fig 4). The final map comprised of all 380 markers mapped onto 20 linkage groups with a total length of 2559.66 cM and a LOD score of 3.5. Segregation distortion analysis revealed that majority of the markers segregating in the population followed a Mendelian segregation pattern of either an F1 (3:1) or F2 (1:2:1). The high proportion of segregation distortion among the markers did not affect their assignment to linkage groups as some of the highly distorted markers were mapped on linkage groups alongside those that were not distorted. Linkage group 1 recorded the largest number of markers comprising of 97 markers with an average marker interval of 3.2cM. Linkage group 20 was the shortest with five markers (Fig 4).

**Fig. 4.**
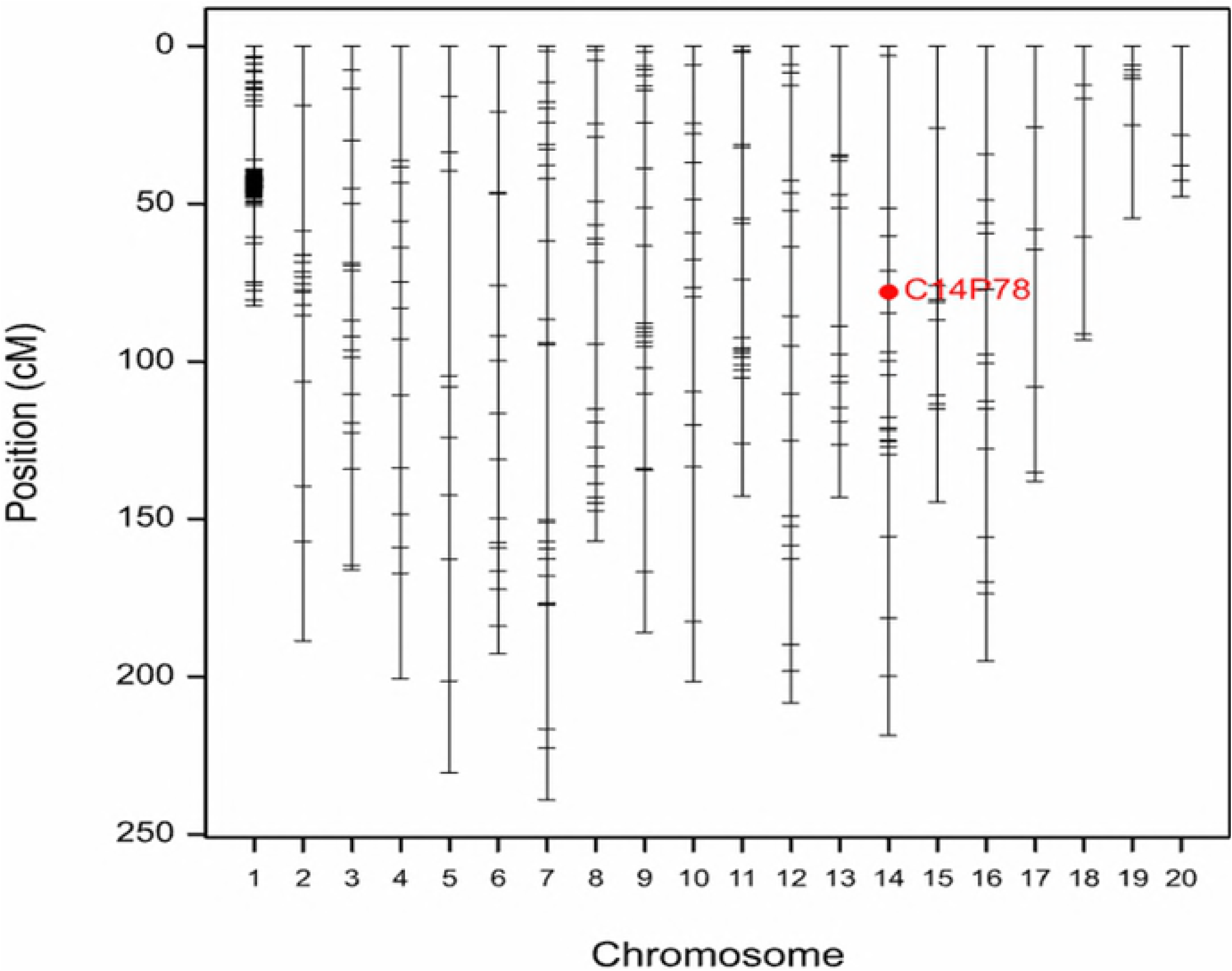
Linkage map consisting of 20 linkage groups constructed from 380 EST-SSR markers. Linkage map constructed based on genotypic data on two parents and 94 progenies.

### QTL Analysis

The *C. gloeosporioides* isolate used in the present study was highly virulent, but had differing responses on both parents and the progenies showing segregation pattern ranging from high resistance with no visual symptoms (average disease severity score of 1.5) to high susceptibility (average disease severity score of 4.5) over three years. The distribution of disease scores for individual three years and averaged over three years for 94 segregating progenies and their parents is represented as a frequency histogram in Figure 3. The distribution of disease severity was skewed towards resistance with majority of the progenies falling under resistance or moderately resistance category (Fig. 3). QTL analysis was carried out using disease severity score data collected during individual years and averaged over three years and SSR data on 94 progenies. Based on simple interval mapping and LOD score of >3.5, a significant QTL was observed on LG 14 at position 77.94 cM (between 71.12 – 84.76 cM) explaining 68.94% of total phenotypic variation (Fig 5). The QTL position was then checked for each year wherein no significant QTL was observed when LOD score was 3.5 for 2011. However, with a lower LOD score of 3.4, a significant QTL was observed on LG 14 (Fig. 6). Similarly, for 2012, one significant QTL was observed on LG 14 with LOD score of 3.8 at position of 77.94 cM (between 71.85 – 84.03 cM) and explaining the phenotypic variation of 75.61%. For 2013, two significant QTL was observed when LOD was greater than 3.5 on LG 12 and LG 14 (Fig 6). The QTL on LG14 was consistently and significantly associated with anthracnose resistance with high LOD score (>3.5) in two out of three years and average score data. This QTL was consistently found at position 77.94 cM falling between 71.12 – 84.76 cM (Table 2). However, if the LOD score was reduced to >2.0, a larger number of QTL (s) were observed across individual years and average score data (Fig 6). Additional QTLs were observed on LG 9, 12 and 17 for 2011; while QTL (s) on LG1, 2, 4, 7 and 11 for 2012; and QTL (s) on LG 9, 11, 12 and 16 for 2013. For average over three years, an additional QTL on LG12 was observed at LOD score of >2.5 (Fig 6; Table 2).

**Fig. 5.**
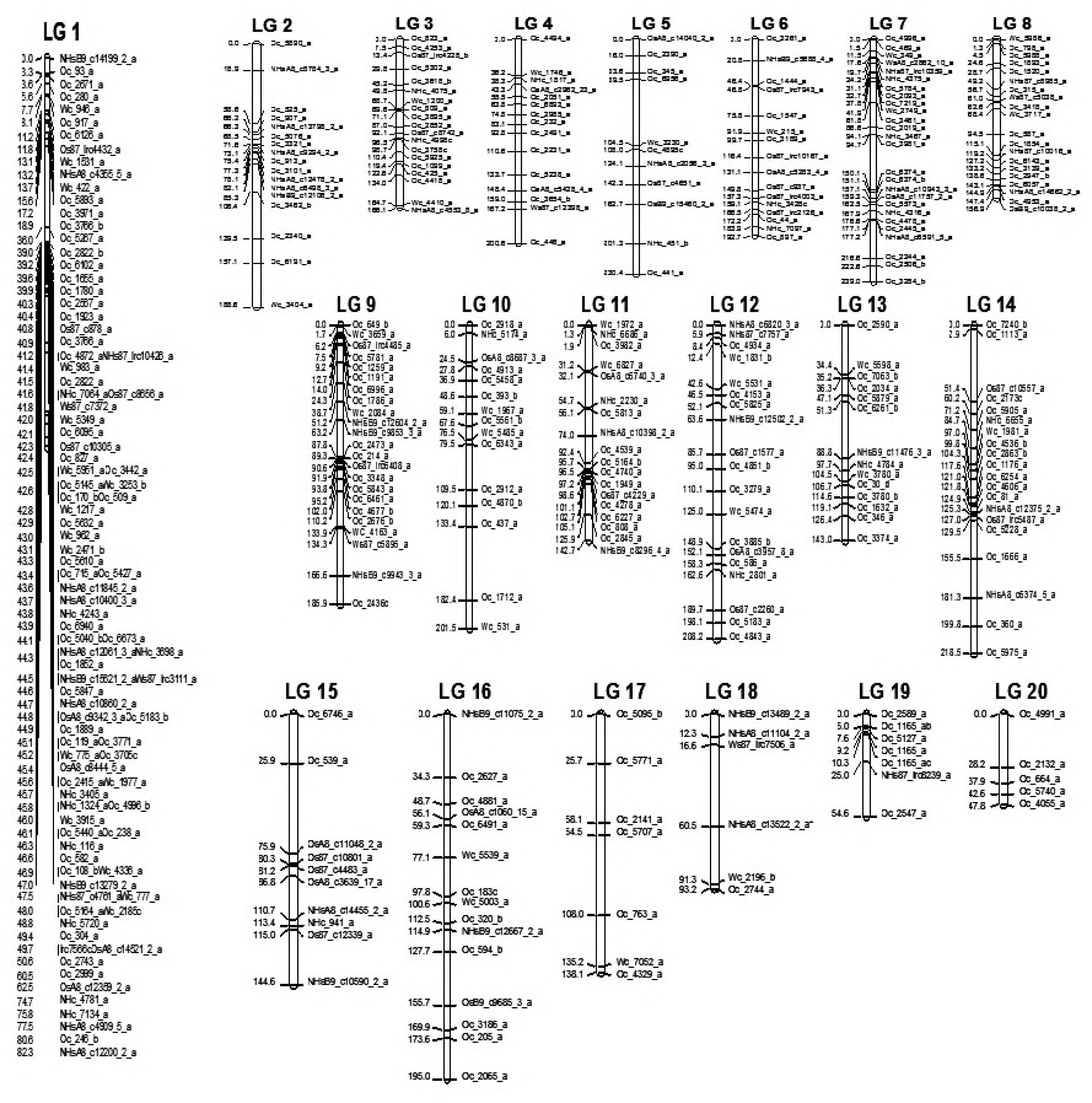
QTL for anthracnose disease based on average score across three years and SSR data with a LOD score of >3.5 using simple interval mapping (SIM) QTL mapping based on EST-SSR data on 94 progenies and mean phenotypic data for anthracnose disease over three years.

**Fig. 6.**
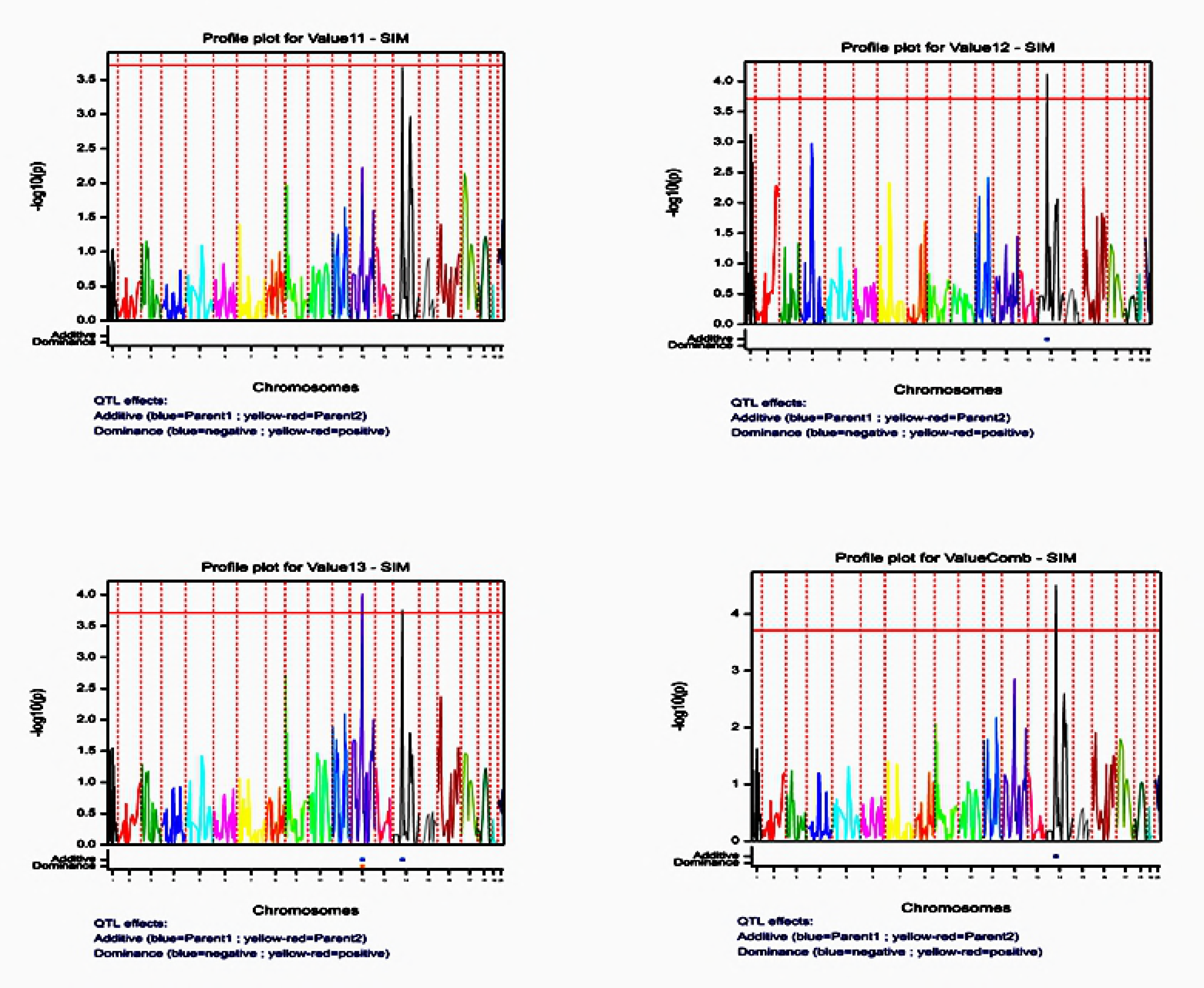
Additional QTL (s) for each individual year and average across three years based on different LOD scores. Additional QTL (s) based on EST-SSR data on 94 progenies and anthracnose disease data for individual years and mean data across all the years at a lower LOD score.

## Discussion

The availability of a saturated genetic linkage map is a valuable tool for genetic research and molecular breeding. The first genetic map of *D. alata* was generated by Mignouna et al [35] using AFLP markers in an intra-specific F1 cross between TDa 95/00328 as female parent and TDa 87/01091 as male parent. Later, Petro et al [19] reported the establishment of an intra-specific genetic map of *D. alata* by crossing two diploid breeding lines, Boutou (female resistant parent) and Pyramide (susceptible male parent) from the Caribbean. The resulting map included 523 polymorphic markers from 26 AFLP primer combinations and covered a total length of 1538 cM including 20 linkage groups. The authors estimated the total length of yam genome to be 1923 cM, and covered over 80% of the genome. In the present study, the linkage map was developed using EST-SSRs that spanned a total length of 2559.66 cM, thus indicating that *D. alata* genome is much longer and whole genome reference sequencing will provide a better understanding of the actual genome size of this crop. The genetic map produced in the present study is moderately saturated with random distribution of EST-SSRs across different linkage groups and it is expected that with the availability of more markers (SSRs or SNPs), a consensus map can be generated with appropriate positioning of markers in each linkage group. The expected genome size of *D. alata* is 620 Mbp with the estimated physical distance per map unit as 620Mbp/2559.66 cM = 242 kb per cM, making map-based gene cloning or QTL analysis as a feasible strategy in water yam. A high level of segregation distortion was observed in this population, which may be due to small size of the mapping population (94 individuals) or genotyping errors or biological factors such as gametic and/or zygotic selection or it could be due to physiological and environmental factors [39, 40, 41]. High level of distortion was also observed by Petro et al [19] while constructing the linkage map from an intraspecific F1 population in *D. alata*.

Both parents used were infected by Kog01R1, although the severity of symptoms was much higher on TDa 95-310 (highly susceptible parent) than on TDa 95/00328 (moderately resistant parent) [14]. The segregation of 94 progenies for anthracnose response showed a distribution from highly resistance to highly susceptible (Fig. 3) although it was skewed towards resistance with more number of progenies falling within moderately resistant and resistant category. This is in support of the findings of Mignouna et al [20] and Petro et al [19] which described that resistance to anthracnose disease is dominantly and quantitatively inherited. The results from the present study clearly demonstrated that the resistance for anthracnose disease in *D. alata* is under polygenic control. Research studies involving host-pathogen systems have showed that susceptible parents do substantially contribute to increasing resistance in the progenies to diseases due to transgressive segregation [42, 43, 44].

This study identified consistent QTL (s) to most virulent isolate of *C. gloeosporioides*, however, the resistance of *D. alata* to anthracnose disease could be isolate-specific or non-specific [20]. There is a need to use combination of two or more isolates to validate the isolate-specificity and non-specificity for resistance, and to understand the isolate-isolate interaction and host-isolate interaction causing anthracnose disease thus throwing light on the existence of the minor-gene-for-minor-gene interaction for quantitative resistance as postulated by Parlevliet and Zadoks [42]. This was based on findings in partially resistant barley lines to *Puccinia hordei* wherein small but significant host-isolate interactions have been reported [43]. Use of QTL-approach to identify isolate specificity for quantitative disease resistance has been reported in several studies [44, 45, 46, 47]. So far, two studies have been undertaken for mapping QTLs controlling resistance or partial resistance to anthracnose disease in water yam [3, 35, 19]. Both studies used AFLP maps and detected one QTL [3, 35] and nine QTLs [19] for partial resistance to anthracnose disease with low to moderate phenotypic variance ranging from 71-84%. However, the non-availability of SSR markers and common linkage group nomenclature in both maps makes it difficult to compare the location of detected QTLs in these two studies. In the present study, we use the same resistant female parent, TDa 95/00328 as used by Mignouna et al [3, 35] and EST-SSRs to develop the linkage map and identify the QTL (s) for anthracnose resistance. We detected one consistent QTL on linkage group 14 across different years of evaluation and average over years that explained about 68.94% of the phenotypic variance in the mapping population progenies, indicating the presence of a probable major QTL in the region. However, this QTL is based on a low number of segregating population (94 progenies) derived from a cross between moderately-resistant and susceptible parents, which will need to be further validated in other mapping populations including highly resistant and susceptible parental genotypes. Similar study can also be carried out using advance technology such as next generation sequencing techniques to validate and develop SNP-chip for marker assisted selection especially at early generation.

## Conclusions

The linkage map obtained in this study showed a good coverage of the probable genome size of water yam, thus allowing for QTL detection and marker-based selection in the breeding activities. This is the first study to report the use of EST-SSRs to generate the linkage map and detection of QTL (s) using the most virulent isolate of *C. gloeosporioides* prevalent in West Africa. This has opened new opportunities for validating the QTL (s) and understanding the host-pathogen interaction for anthracnose disease in *D. alata*. Additional markers including SSRs, SNPs, etc. will further elucidate the importance of candidate gene approach for marker-assisted selection in the improvement of this important staple food crop.

## Acknowledgments

The authors greatly acknowledge the USAID-Linkage grant awarded through IITA to Ranjana Bhattacharjee. Authors would like to acknowledge the CGIAR Research Program on Roots, Tubers and Bananas (CRP-RTB) for additional financial support. Authors would like to acknowledge Drs. Antonio Lopez-Montes and Lava Kumar for providing the mapping population and isolates of *C. gloeosporioides* for anthracnose disease screening; Christian Nwadili for phenotypic evaluation in the screenhouse; and laboratory personnels from Saski’s team for EST-SSR genotyping.

